# Functional metagenomics of bark microbial communities from avocado trees (*Persea americana* Mill.) reveals potential for bacterial primary productivity

**DOI:** 10.1101/2020.09.05.284570

**Authors:** Eneas Aguirre-von-Wobeser

**Affiliations:** CONACYT – Centro de Investigación y Desarrollo en Agrobiotecnología Alimentaria, Centro de Investigación y Desarrollo, A.C., Blvd. Sta. Catarina s/n, Col. Santiago Tlapacoya, 42110, San Agustín Tlaxiaca, Hidalgo, Mexico

**Keywords:** Bark microbial communities, Metagenomics, Plant microbiome, Bacteria, *Persea americana* (avocado), Prokaryotic photosynthesis

## Abstract

Bark microbial communities are poorly understood, and information on the metabolic capacities of their inhabitants is lacking. Bark microbial communities share part of their taxonomic composition with soil, but the functional differences and similarities are unknown. By comparing bark microbial communities of avocado trees (*Persea americana*, Mill.) with rhizospheric soil, functional processes relevant to the bark environment were identified. DNA from bark and soil communities was extracted from the same trees, and shotgun metagenomics sequencing was performed using nextSeq technology. Genes were identified by BLAST methods, and functional annotation was performed with KEGG databases as a reference. Bacterial oxygenic and anoxygenic photosynthesis genes were highly abundant in bark as compared to soil. Furthermore, increased presence of nitrogenase genes suggests a potential for nitrogen fixation. Genes for methanol utilization were abundant in bark, but no evidence of methane utilization potential was observed. Bark microbial communities have the genetic information for potential primary productivity, which might contribute to microbial growth independent of plant-derived carbon substrates.

## Introduction

Plants live in association with diverse communities of microorganisms, many of which have beneficial or detrimental interactions with their hosts (Hassani *et al*., 2018), and others inhabiting plant tissues as commensals. The plant microbiome is not limited to the rhizosphere, as consistent taxonomic groups have been observed in aerial plant tissues as well (Lambais *et al*., 2014; Coleman-Derr *et al*. 2015; Aguirre-von-Wobeser *et al*., 2020; Wu *et al*., 2020). The complexity of microbial communities on aboveground plant tissues has been established by ample literature on leaf microbiology (Stone *et al*., 2018), where microorganisms are commonly found aggregated on protected grooves (Vorholt, 2012).

Plant surfaces are considered a challenging environment for microbial life (Stone *et al*., 2018), with high exposure to ultraviolet radiation (UV), low availability of water, nutrient scarcity and the presence of antimicrobial compounds (Vorholt, 2012). However, conclusions drawn for studies on leaves cannot be directly extrapolated to the bark environment. For instance, inner stems, including the trunk of trees, can be shaded by the canopy from direct UV and high photosynthtic active radiation (Muñoz-García *et al*., 2014), which could decrease DNA damage and oxidative stress. Although water scarcity could be present, its extent is not clear, as bark can store considerable amounts of water (Rosell *et al*., 2014) and retain humidity (Mehltreter *et al*., 2005), which could also be aided by canopy shading. On the other hand, the chemical environment in bark could be at least as challenging as the leafs (Tanase *et al*., 2019). *Persea americana* (Mill.) bark, for example, is known to contain compounds with bactericidal activity (Akinpelu *et al*., 2014).

Adaptations to the potential stresses found on plant surfaces have been observed on the phylosphere (Vorholt, 2012; Stone *et al*., 2018), which might be relevant on bark as well. To cope with desiccation stress, bacteria can form biofilms, which can retain water (Roberson and Firestone, 1992; Gorbushina, 2007; Kakumanu *et al*., 2013). The effects of UV radiation exposure can alleviated with photoprotective pigments (Jacobs *et al*., 2005; Vorholt, 2012) and enzymes or compounds to combat oxidative stress, like catalase and superoxide dismutase or tocopherols (Sakuragi *et al*., 2006).

Nutrient scarcity is also considered a stressful condition on plant surfaces, including nitrogen limitation (Vorholt, 2012; Stone *et al*., 2018). Nitrogenous compounds present in bark include amino acids, proteins, ammonium and nitrate (O’Kennedy and Titus, 1979; Gowda *et al*., 1988; El-Hamalawi and Menge, 1994; Wolterbeek *et al*., 1996; Spangenberg *et al*., 2002). Nitrogen fixation can also provide nitrogen for some microorganisms in plant surfaces, including bark (Yatazawa *et al*., 1983).

Carbon limitation can also be severe in plant aerial surfaces. While it is recognized that carbon availability is low on leaf surfaces (Vorholt, 2012; Stone *et al*., 2018), there could be more carbon sources to support microbial growth in bark, including complex carbohydrates (Gowda *et al*., 1988). Bacteria able to degrade and utilize complex bark carbohydrates like pectin, xylan, lignin, tannins and phenolic compounds have been reported on decaying wood (Deschamps *et al*., 1980; Deschamps *et al*., 1983), and some have been isolated from living trees (Erakhrumen and Boyi, 2016). Transporters of carbon molecules are prevalent in the phylosphere, further supporting the evidence of utilization of diverse carbon substrates on plant surfaces (Lambais *et al*., 2017). However, the potential of plant aerial surface microbial communities of acquiring carbon by photosynthesis has not been considered in the literature.

Microbial communities of bark have similar phylum-level phylogenetic structures as the phylosphere, where Proteobacteria, Actinobacteria and Bacteroidetes are dominant (Vorholt, 2012; Lambais *et al*., 2014; Leff *et al*., 2015; Arrigoni *et al*., 2018; Vitulo *et al*., 2019; Aguirre-von-Wobeser *et al*., 2020; Arrigoni *et al*., 2020). However, it is unknown if the functional repertoires of these communities have similar properties. In this work, the bark of *P. americana*, was analyzed, to identify metabolic networks present the bacterial component of its microbial communities. Abundant prokaryotic photosynthetic genes for both oxygenic and anoxygenic photosynthesis were found, as well as genes encoding nitrogen fixation enzymes. These results suggest that bark microbial communities have an important potential for primary productivity, which could contribute to sustain their populations in addition to three-generated organic matter.

## Materials and Methods

### Sampling procedures

Samples collection was described previously (Aguirre von Wobeser *et al*., 2020). This study focuses on avocado trees from an orchard located near Malinalco, Mexico (18° 59’ 22.2828’’ N, 99° 29’ 40.4124’’ W). The orchard was selected to compare rhizospheric soil and bark functional potential of microbial communities, as metagenomic DNA reads were available for three bark and three rhizosphere samples from the same trees. A detailed description of the orchard is available (Aguirre-von-Wobeser *et al*., 2020). Briefly the orchard is located at approximately 2000 m above sea level, and is managed with organic practices, with no use of agrochemicals. It was abandoned for 18 years, and recently prunned and cleaned; therefore, it has a long history without agrochemical use. Weeks before sampling, a preparation containing *Trichoderma harzianum* and *Glomus* sp. was added to the soil. The sampled trees were located in the same of the orchard, and were healthy, productive trees. Three sample pairs of bark and rhizosperic soil yielded high quality metagenomic sequences, and were included in the present study.

### DNA extraction, library preparation and sequencing

Total DNA was extracted with a PowerSoil kit as described in Aguirre-von-Wobeser *et al*. (2020). To introduce bark samples to the PowerSoil kit vials, the periderm, which is the outermost layer, was collected using a razor blade and cut into small pieces. Libraries were prepared with a Nextera DNA Flex Library Prep Kit, using a Netxtera DNA CD Indexes kit, following manufacturer’s instructions. Sequencing was performed on a nextSeq sequencer (Illumina, USA) using standard procedures. Raw sequencing data can be downloaded from the NCBI Short Read Archive (BioProject PRJNA656796, accession SUB7881803) and from KBase (https://kbase.us/n/69195/32/).

### Bioinformatic analysis

Avocado host contamination was tested using Bowtie (Langmead *et al*., 2009), and was less than 2.3% for all samples (Aguirre-von-Wobeser *et al*., 2020). To remove low quality nucleotide-calls, sequence reads were processed with Trimmomatic (Bolger *et al*., 2014), using a sliding window of 4 nucleotides and a quality filter of 15 (Phred scale), removing all relevant Illumina sequences, and pair-end reads were joined with vsearch (Rognes *et al*., 2016). To increase the proportion of coding regions, random subsets of 10 million reads per sample were aligned to the bacterial protein sequences in the RefSeq database (O’Leary *et al*., 2016). This was conducted using the BLASTx program run with Diamond (Buchfink *et al*., 2015). The reads which had significant matches to RefSeq were transcribed to amino acids in the appropriate reading frame, and were used for functional annotation with GhostKoala (Kanehisa *et al*., 2016), which uses the KEGG database as a reference (Kanehisa and Goto, 2000).

Reads with matches to the same orthologs in the KEGG database were counted, as a measure of the abundance of the corresponding gene in each sample. Since the abundances for all sample-pairs were linearly correlated along the whole observed abundance range (Supporting Information Fig. S1, see Results section), a simple linear transformation was sufficient for adequate normalization, by dividing the abundances of each sample by their respective sums, and multiplying by the mean of the sums for all samples. To test for differential abundance of the annotated orthologous reads for each gene, student’s t-tests were conducted based on normalized log-transformed abundances. For these tests, only genes present in all samples were considered. The obtained p-values were corrected for multiple testing with False Discovery Rates (Benjamini and Hochberg, 1995) using the package p.adjust from R, and the abundances were considered statistically significant with adjusted p values less than 0.05, *i. e*., 5% of the genes with significantly different abundances were expected to be false positives. The distribution of p-values was adequate for multiple parametric statistical tests (Supporting Information Fig. S2, see Results section) as discussed by Storey and Tibshirani (2003), suggesting that the logarithm-transformed data were amenable to t-tests. In addition to genes with statistically significant differences in abundance, genes which were found in all samples from one group (either bark or soil), but were absent in all samples from the other, were considered as more abundant in the group in which they were observed.

## Results

### Shotgun metagenomics dataset

In this study, the analysis on avocado bark microbial communities is deepened, using a data-set presented before, where the taxonomic composition was discussed in detail (Aguirre-von-Wobeser *et al*., 2020). From this dataset, the three sample-pairs where bark and rhizosphere samples were available for the same tree were selected. Of these, reads per sample ranged from 88 x 10^6^ to 133 x 10^6^ for bark, and from 62 x 10^6^ to 190 x 10^6^ for rhizospheric soil.

Random samples of 10 million reads were aligned to the proteins in the RefSeq database. Of these, 40 to 54% of the reads had matches to bacterial RefSeq proteins, and 8.9 to 12.75% could be annotated with the KEGG database (Supporting Information Table S1). These annotations constitute reads homologous to parts of genes; however, for brevity they are referred to as “genes” in this manuscript, but caution should be in place, as they do not constitute solid proof of the presence of functional genes. Read counts for all the annotated genes showed extremely high correlations within sample groups (bark and soil), showing that the dataset had high consistency between independent samples (Supporting Information Fig. S1). Furthermore, samples taken from the two groups had high correlations between them, suggesting that the bacterial populations of these microbial communities were not too divergent, which is consistent with our previous taxonomic analysis (Aguirre-von-Wobeser *et al*., 2020). Nevertheless, 602 genes had significant differences in their abundances (Supporting Information Fig. S2) between both environments (312 higher in bark and 290 higher in rhizospheric soil). Furthermore, 102 were found only the bark environment (in all three samples), and 143 were found only in rhizospheric soil (in all three samples). These differences likely represent adaptive functions for the environmental conditions faced in bark and rhizospheric soil, respectively. The results for all statistical tests and the read counts in each of the samples can be found at Supporting Information File S1.

### Central metabolism and housekeeping functions

Genes for the main routes of central metabolism were present in the bark and rhizospheric soil environments, including glycolysis, the Krebs cycle, the pentose phosphate cycle, amino acid metabolism and lipid metabolism (Supporting Information File S2). A complete set of orthologs of genes for housekeeping cellular functions, including DNA replication and protein synthesis, was also found in both environments.

#### Photosynthetic light reactions and pigments

Genes related to prokaryotic photosynthesis were consistently more abundant in bark, as compared to rhizospheric soil microbial communities. For example, many genes for the synthesis of chlorophyll *a*, mostly from *Cyanobacteria* and *Alphaproteobacteria*, were significantly more abundant in bark as compared to rhizospheric soil (Fig. 1a), suggesting a major roll of oxygenic photosynthesis in bark microbial communities. Furthermore, genes for key enzymes in the production of bacteriochlorophyll *a* and *b* were also enriched in bark communities, including *bchY*, encoding for a 3,8-divinyl chlorophyllide a/chlorophyllide a reductase, which is the first dedicated enzyme in the pathway of bacteriochlorophyll synthesis (Figs. 1a and 2a). Orthologs of geranylgeranyl diphosphate/geranylgeranyl-bacteriochlorophyllide *a* reductase, encoded by c*hlP*, from *Cyanobacteria*, *Proteobacteria* and *Gemmatimonadetes* were almost exclusively found in bark, further suggesting that bacteriochlorophyll is also produced in this environment (Figs. 1a and 2b).

**Figure 1.**
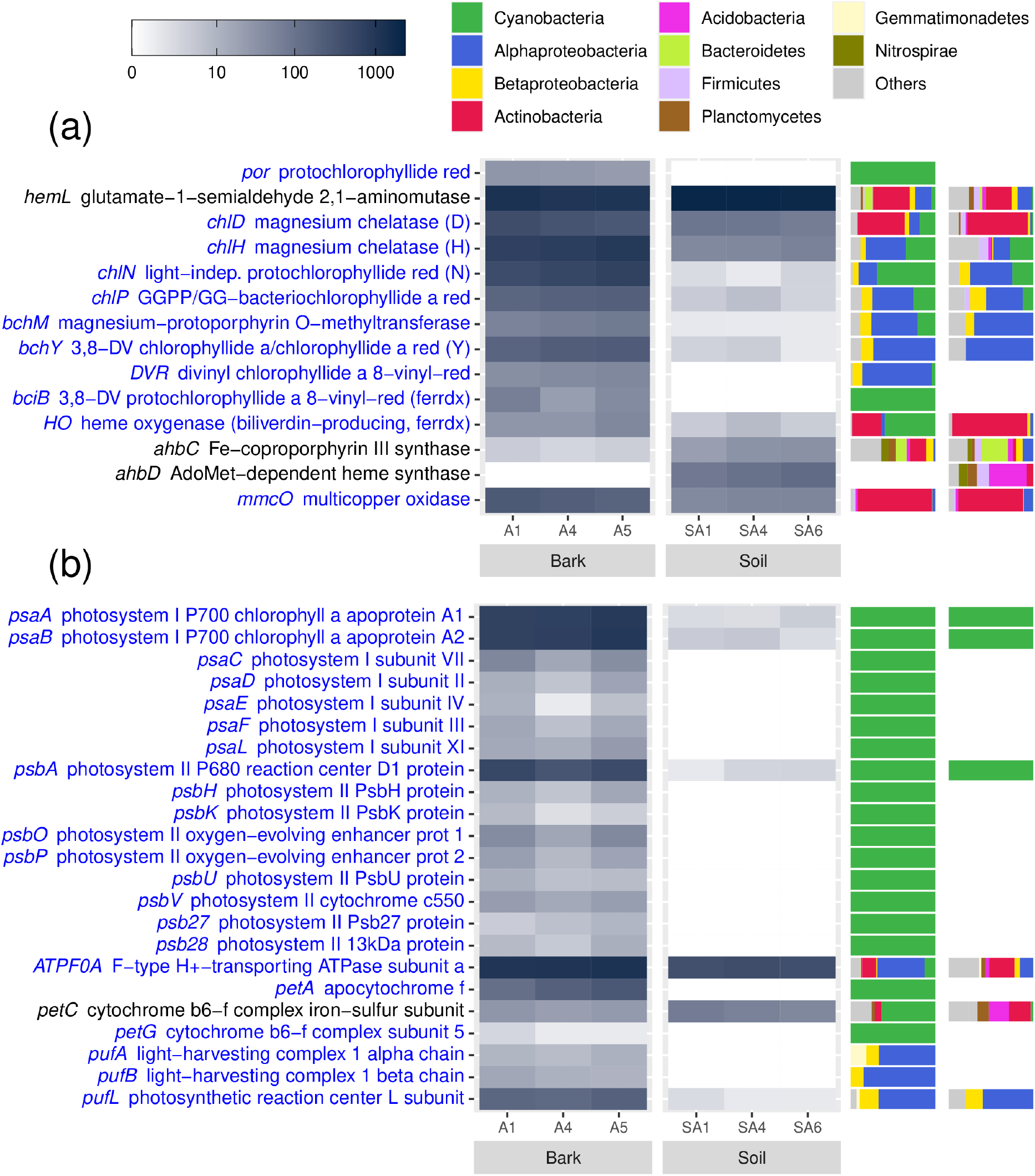
*Photosynthesis-related genes with different abundance in Persea americana* (Mill.) *bark and rhizospheric soil*. (a) Chlorophyll and bacteriochlorophyll synthesis genes. (b) Photosynthetic apparatus genes. Genes shown had significant differences between bark and rhizospheric soil (adjusted p < 0.05). Blue font in row labels indicates a higher abundance in bark. The scale shows the number of reads identified for each group, from a randomly selected set of 10,000 reads for each biological sample.

**Figure 2.**
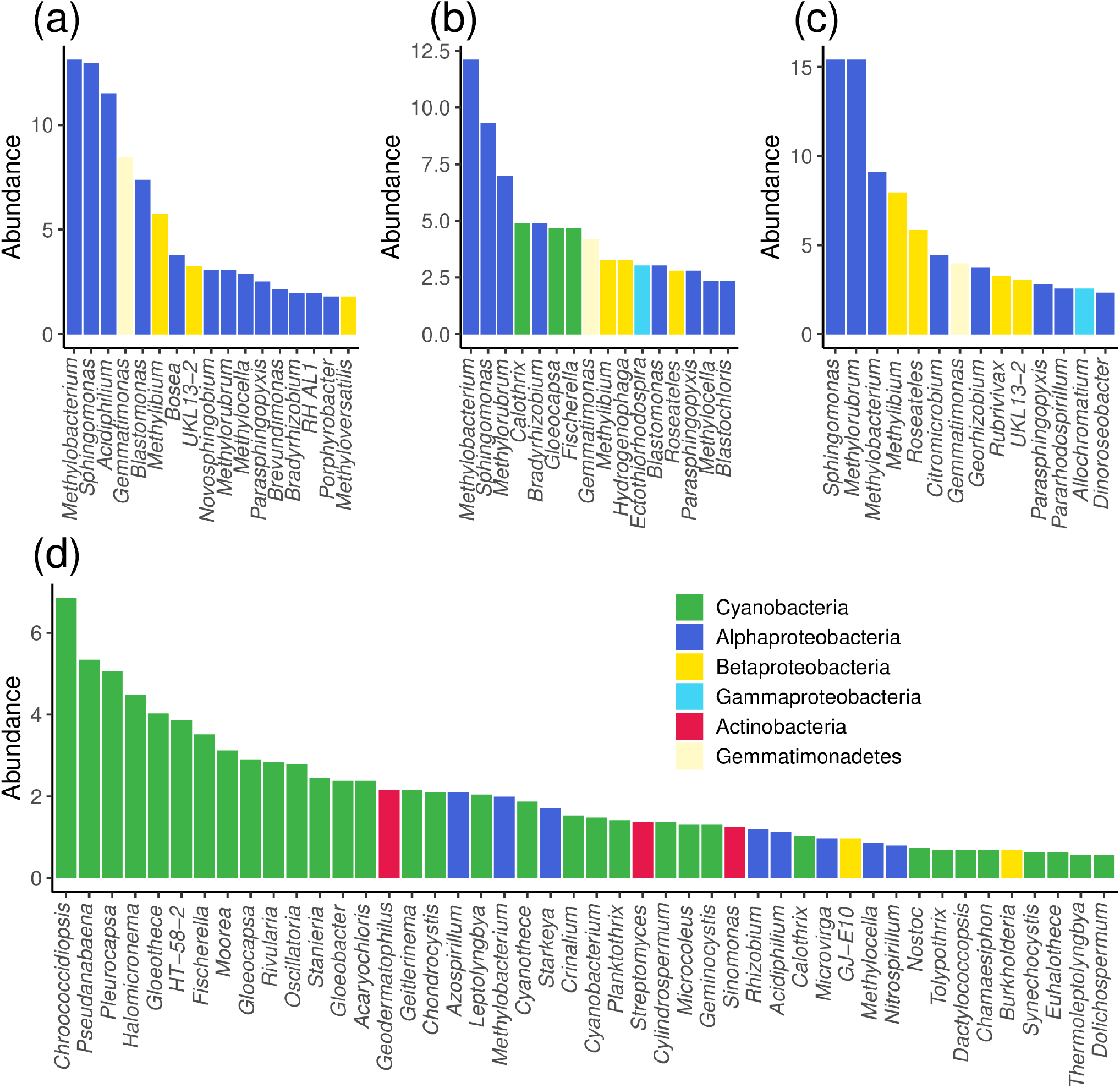
*Taxonomic annotation of reads from selected photosynthesis-related genes found in Persea americana* (Mill.) *bark*. (*a) bchY, 3,8-divinyl chlorophyllide a/chlorophyllide a reductase subunit Y (b) chlP, geranylgeranyl diphosphate/geranylgeranyl-bacteriochlorophyllide a reductase (c) pufL*, photosynthetic reaction center L subunit *(d) rbcL, ribulose-bisphosphate carboxylase large chain. Bars represent the percentage of reads belonging to each genus, from all samples combined, relative to the total number of reads detected for the corresponding gene*.

Some genes with roles in the synthesis pathways of chlorophylls, like *hemL, mmcO, HO* and *ChlD*, were common in bark and rhizospheric soil Actinobacteria; however, these genes are not specific to chlorophyll synthesis, and are likely involved in other biochemical processes in this taxonomic group. *HemL* is very common in bacteria, as it is essential for the biosynthesis of all tetrapyrroles, and *HO* is generally used to degrade heme to obtain iron (Dailey *et al*., 2017). *MmcO* genes produce enzymes which oxidize a large variety of substrates, including reactive oxygen species, different metals and complex organic molecules (Kinkar *et al*., 2019). *ChlD* genes are involved in cobalamin synthesis in many Actinobacteria (Antonov, 2020).

Genes for the production of phycobilins, the chromophores of phycobiliproteins, were exclusively found in bark (Supporting Information Fig. S3a), which is consistent with the higher abundance of cyanobacteria in these microbial communities as compared to rhizospheric soil (Aguirre-von-Wobeser *et al*., 2020). Similarly, genes for the phycobiliproteins, all from cyanobacteria, were only observed in bark (Supporting Information Fig. S3b).

Genes encoding for cyanobacterial (plant-like) photosystems I and II were also more abundant in bark than in rhizospheric soil (Fig. 1b), indicating a higher prevalence of the photosynthetic electron transport chain apparatus in this environment. Also, genes for an anoxygenic photosystem, pufA, pufB and pufL were more abundant in bark, mostly from Alphaproteobacteria, but also from Betaproteobacteria and Gemmatimodadetes (Figs. 1b and 2c). More taxonomic diversity was observed for ATP-ase and cytochrome genes, but these may be related to respiratory electron transport chains.

#### Carbon fixation

Genes for all the enzymes participating in the Calvin-Benson cycle were detected in both rhizospheric soil and bark samples, suggesting the potential for microbial carbon fixation in both environments (Supporting Information Table S1). However, genes for some of these enzymes were more abundant in bark. Notably, the key enzyme Ribulose-1,5-bisphosphate carboxylase/oxygenase (RuBisCO), which is utilized in both oxygenic and anoxygenic photosynthesis for carbon fixation, was more abundant in bark than in rhizospheric soil microbial communities (Fig 2d, Supporting Information Fig. S4). Reads mapping to this enzyme were found in Cyanobacteria, Proteobacteria (alpha and beta) and Actinobacteria. Although some of these genes in Actinobacteria are likely to be RuBISCO-like enzymes, also known as type IV RuBISCO, which catalyze different reactions (Selesi *et al*., 2005), in the case of *Geodermatophilus*, photosynthetic carbon fixation capability has been suggested (Sghaier *et al*., 2016).

Genes for alternative carbon fixation pathways found in certain groups of prokaryotes were found in both bark and rhizospheric soil. Nevertheless, these pathways were more complete in rhizospheric soil than in bark. From the genes detected in bark overall (including those not different in their abundance compared to soil), it can be inferred that carbon fixation in this community is mainly conducted through the Calvin Cycle. It is also possible that the Arnon-Buchanan cycle (also known as the Reductive Citric Acid Cycle) is present, as some versions of which had all their enzymes present in bark (Supporting Information Table S2).

#### Methylotrophy

Both bark and rhizospheric soil had almost complete methane metabolism pathways (Supporting Information File S2), suggesting that methylotrophy is important in both environments. However, no methane monooxygenase genes, which are essential for methane utilization, were observed in bark. In soil, two subunits annotated as methane/ammonia monooxygenase were detected, but they seem to be involved in nitrification, rather than methanotrophy, as they belong to groups like Nitrospirae and Actinobacteria (Fig. 3a). As expected, many Alphaproteobacteria from bark, including *Methylobacterium, Methylorubrum, Methylocella* and *Nitrospirillum* had a methanol dehydrogenase gene (*xoxF*), as well as other Proteobacteria, and the Gemmatimonadetes *Gemmatirosa* (Fig. 3b). However, the abundance of *xoxF* was lower in bark than in rhizospheric soil (Fig. 3a). Another methanol dehydrogenase (*mdo*), belonging to different Actinobacteria (Fig. 3c), was more abundant in bark. The specific electron acceptor cytochrome c-L (*mxaG*), was only detected for *Methylobacterium* from bark.

**Figure 3.**
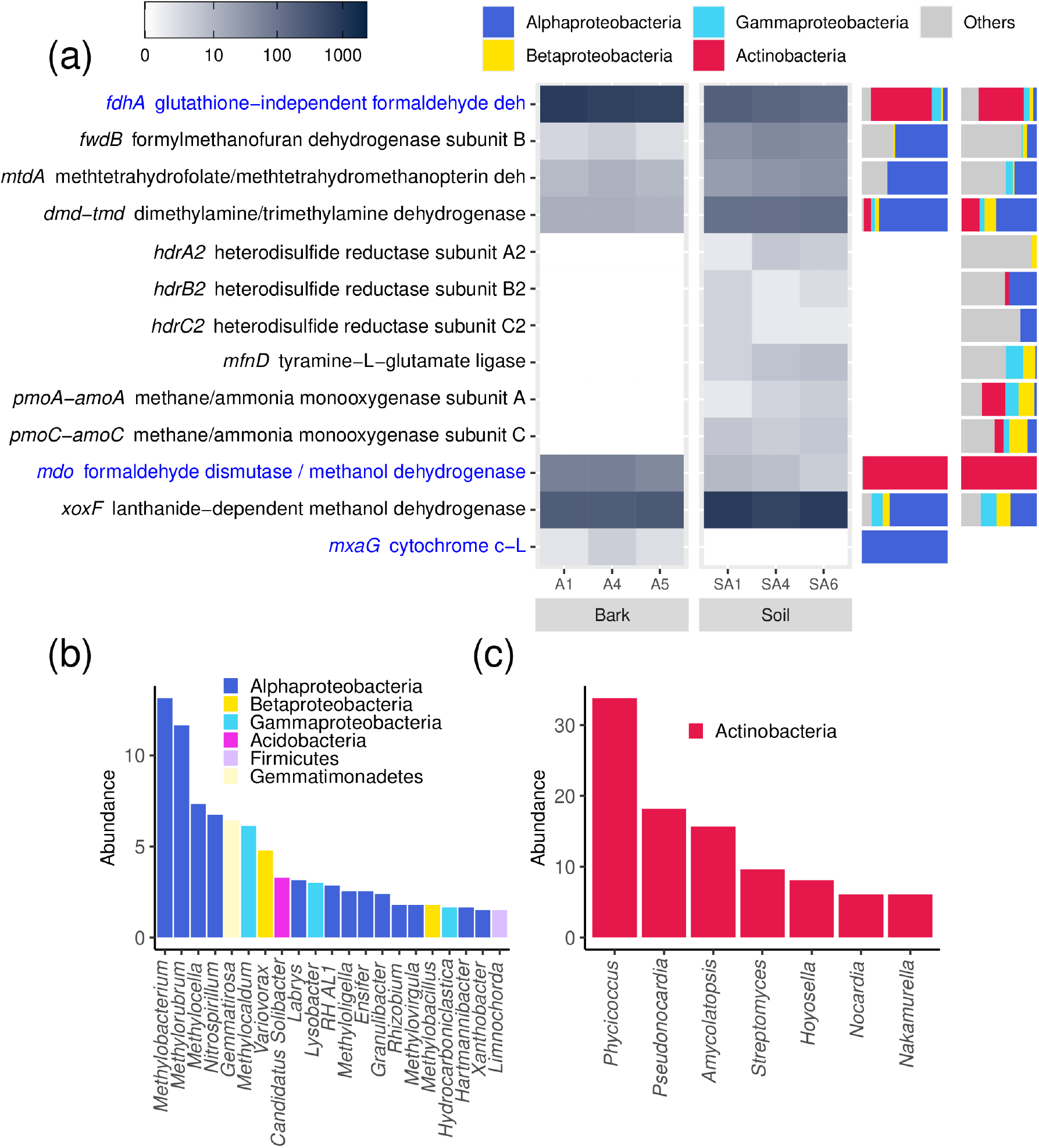
Methylotrophy-related genes with different abundance in *Persea americana* (Mill.) bark and rhizospheric soil. (a) Genes with significant differences between bark and rhizospheric soil (adjusted p < 0.05). Blue font in row labels indicates a higher abundance in bark. The scale shows the number of reads identified for each group, from a randomly selected set of 10,000 reads for each biological sample. (b) Taxonomic annotation of reads for gene *xox*, flanthanide-dependent methanol dehydrogenase, found in bark. (c) Taxonomic annotation of reads for *mdo*, formaldehyde dismutase / methanol dehydrogenase, found in bark.

#### Nitrogen metabolism

Both rhizospheric soil and bark had extensive repertoires of nitrogen metabolism genes (Supporting Information Table S1). However, nitrogenease genes were more abundant in bark than in rhizospheric soil, suggesting nitrogen fixation is more prevalent in this environment (Fig. 4a). Most nitrogenase reads detected were from heterocyst-forming *Cyanobacteria* (Figs. 4b and 4c). Another important process of the nitrogen cycle, denitrification, was apparently more relevant in rhizospheric soil, as genes for two enzymes of this process were exclusively found in rhizospheric soil (Fig. 4a).

**Figure 4.**
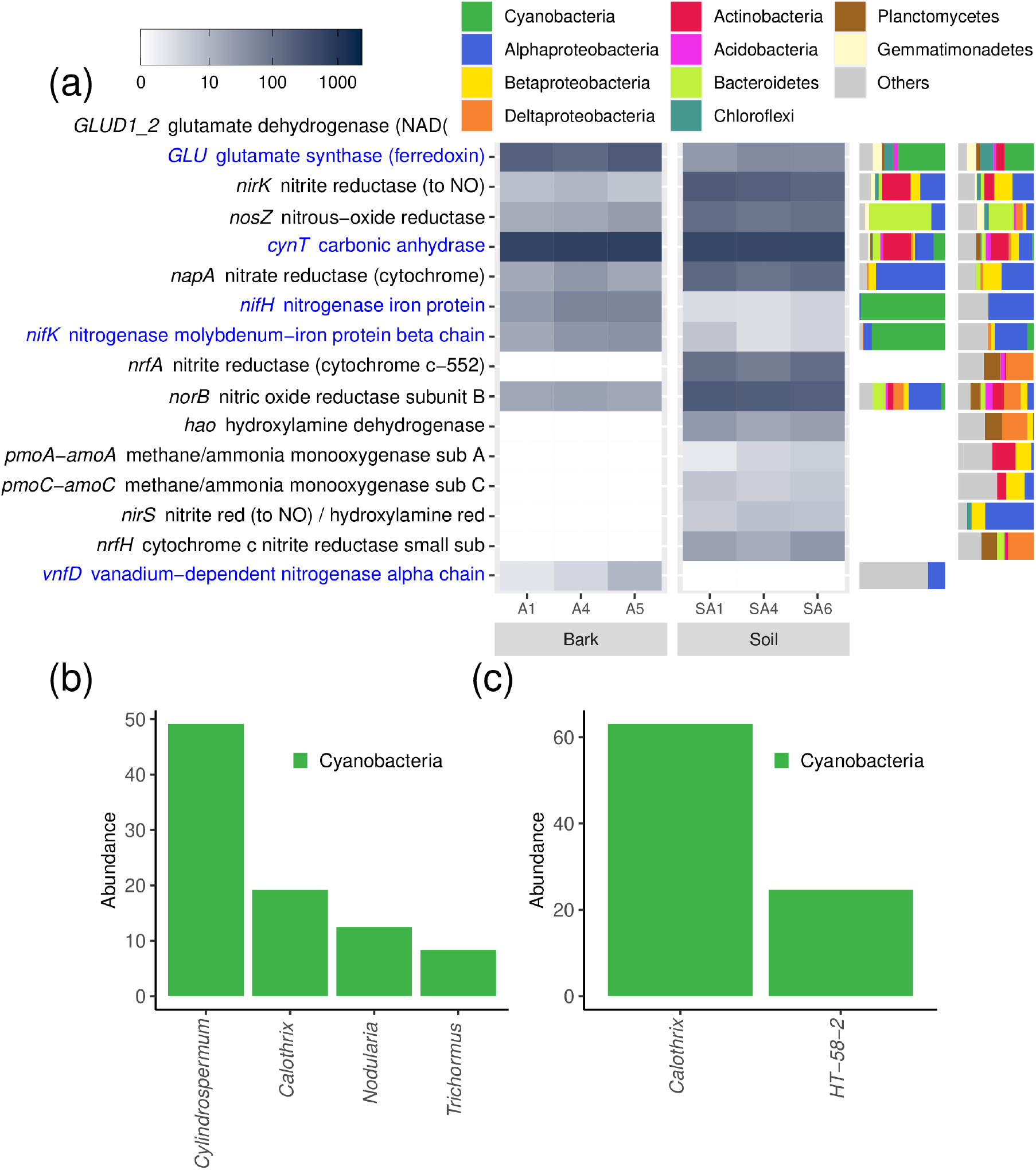
Nitrogen metabolism-related genes with different abundance in *Persea americana* (Mill.) bark and rhizospheric soil. (a) Genes with significant differences between bark and rhizospheric soil (adjusted p < 0.05). Blue font in row labels indicates a higher abundance in bark. The scale shows the number of reads identified for each group, from a randomly selected set of 10,000 reads for each biological sample. (b) Taxonomic annotation of reads for gene *nifH*, nitrogenase iron protein NifH, found in bark. (c) Taxonomic annotation of reads for *nifK*, nitrogenase molybdenum-iron protein beta chain, found in bark.

#### Carotenoid pigments and oxidative stress protection

Genes involved in the synthesis of several caroteinoids were more abundant in bark than in rhizospheric soil (Fig. 5a). These included genes involved in the synthesis of Lycopene, alpha-Carotene, beta-Carotene, Isorenieratene and others. These genes were associated with different taxonomic groups, including Cyanobacteria, Actinobacteria, Proteobacteria, Bacteroidetes and Planctomycetes.

**Figure 5.**
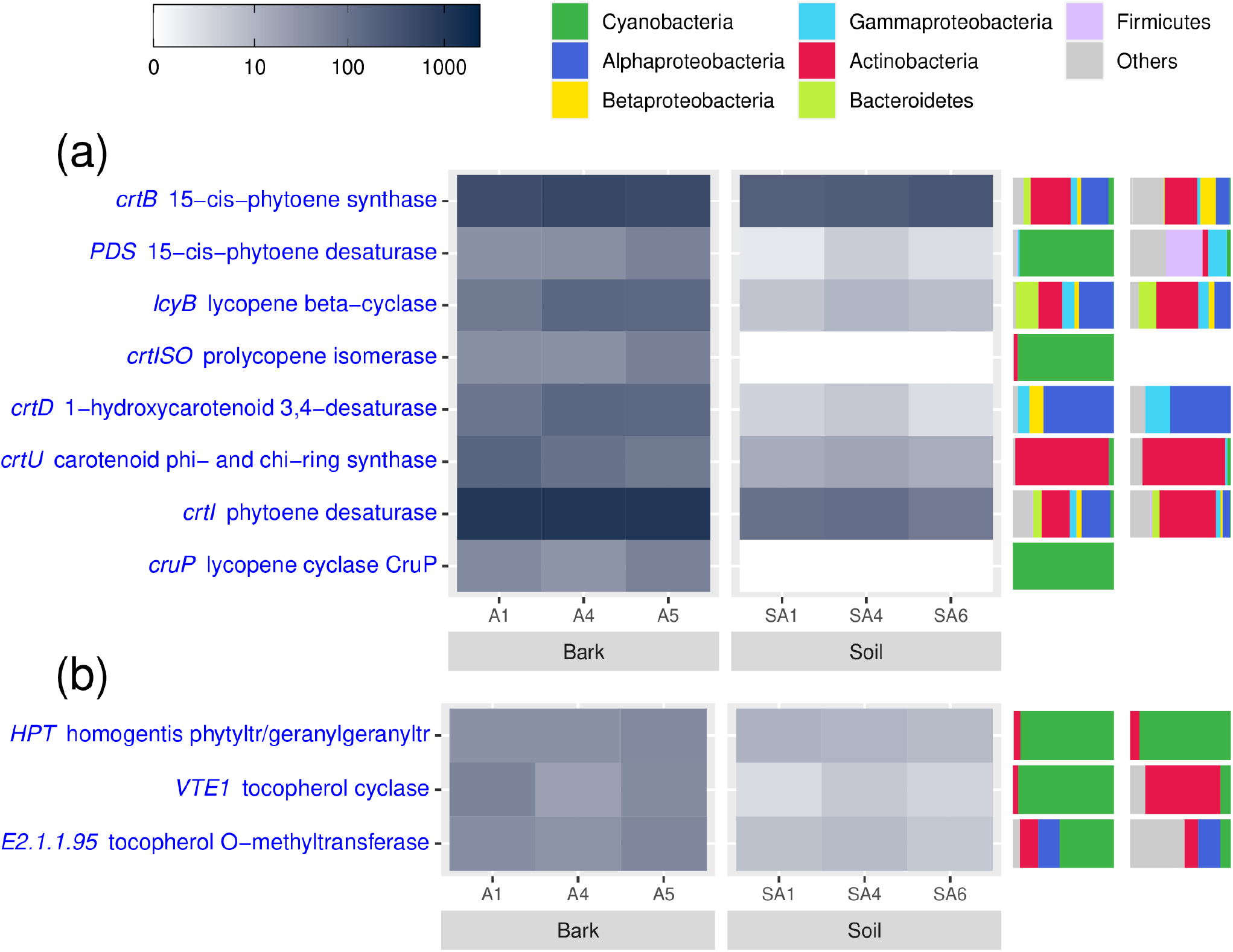
Carotenoids and tocopherol synthesis related genes with different abundance in *Persea americana* (Mill.) bark and rhizospheric soil. Genes shown had significant differences between bark and rhizospheric soil (adjusted p < 0.05). Blue font in row labels indicates a higher abundance in bark. The scale shows the number of reads identified for each group, from a randomly selected set of 10,000 reads for each biological sample.

Genes for the synthesis of tocopherol, an important antioxidant, were also more abundant in bark than in rhizospheric soil, mostly from Cyanobacteria (Fig. 5b).

#### ABC transporters and organic matter degradation

The presence of genes for ABC transporters was markedly different between bark and rhizospheric soil microbial communities (Fig. 6). There were many genes for sugar and sugar alcohol transport from Actinobacteria and Alphaproteobacteria in bark communities, including xylose, frutose, galactofuranose, cellobiose, xylitol and sorbitol/mannitol. Other ABC transporters with more abundance in bark were for transport of different aminoacids, mainly from Actinobacteria and Alphaproteobacteria, but also from Cyanobacteria. Genes for bicarbonate transport from Cyanobacteria, which are part of their Carbon-Concentrating Mechanism, were also more abundant in bark, as compared to rhizospheric soil.

**Figure 6.**
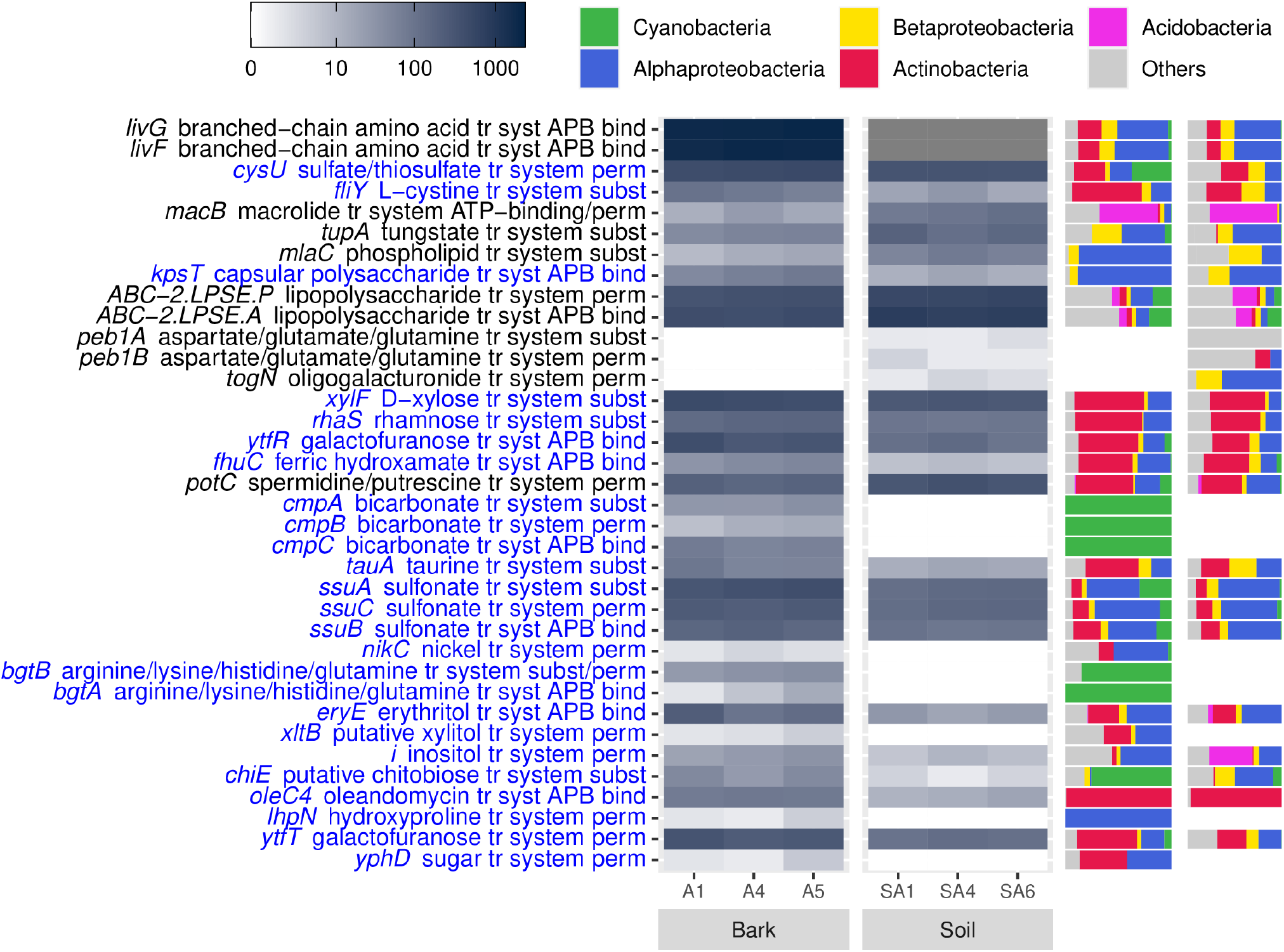
ABC-transporter genes with different abundance in *Persea americana* (Mill.) bark and rhizospheric soil. Genes shown had significant differences between bark and rhizospheric soil (adjusted p < 0.05). Blue font in row labels indicates a higher abundance in bark. The scale shows the number of reads identified for each group, from a randomly selected set of 10,000 reads for each biological sample.

There were several genes with statistically significant different abundance between bark and rhizospheric soil in pathways for sugar transformations (Supporting Information Fig. S6). However, as these pathways can use the same enzymes for catabolic and anabolic reactions, it is difficult to assess whether they increase the potential of sugar utilization by bark or soil bacteria.

Both bark and rhizosphere bacterial communities had many genes for the degradation of complex carbon molecules, including many aromatic compounds, carbohydrates, lipids, amino acids, etc. (Supporting Information File S2). However, very few of these genes had significantly different abundances, suggesting that complex organic carbon degradation and utilization is not specifically increased in either environment.

#### Bacterial interactions, motility and biofilm formation

Genes for exopolysaccharide and surfactant production, quorum sensing, flagellar motility and antimicrobial substance production were widespread in both bark and rhizospheric soil (Supporting Information Table S1), with few significant differences between soil and bark samples.

## Discussion

Despite the widespread distribution of trees in the world (Crowther *et al*., 2015), bark as a microbial habitat has been largely neglected in the literature. The importance of different plant tissues, other than roots and leaves, as distinct microbial environments is gaining recognition in recent years (Ottesen *et al*., 2013; Hacquard and Schadt, 2015; Coleman-Derr *et al*. 2016). In a previous article, we presented the taxonomic composition of avocado bark microbial communities (Aguirre-von-Wobeser *et al*., 2020), showing a high consistency of dominant taxa, even in trees from two different geographic locations. To our knowledge, the present report is the first functional metagenomic analysis of bark microbial communities performed on any woody plant, providing insights on the repertoire of potential functions encoded in their DNA. Cheng *et al*. (2018) analyzed the microbial communities of bark beetle galleries in pine trees using metagenomics, which reside in the phloem, a living tissue, located inwards of the periderm. The present study, in contrast, focuses on the external layer of bark, and was targeted at epiphytes, and possibly includes endophytes residing close to the surface of the trunk (main stem of the tree).

Some biochemical processes are central to microbial cell functioning, and are present in virtually any microorganism, or are very widespread, and should be found in any complex community. The presence of these pathways was verified in our results, to confirm that the functional repertoire identified was comprehensive. Complete pathways for glycolysis, the Krebs cycle, the pentose phosphate cycle, amino acid biosynthesis, lipid biosynthesis and degradation, nucleotide synthesis, gene expression and protein synthesis, and others, were found (Supporting File 2). This suggests that our sequencing effort was thorough, providing a complete picture of the bacterial metabolism in these microbial communities. Limitations of our study must however be considered, as the presence of DNA in a community does not ensure that it is located in viable cells, or that it is transcribed into functional proteins. Furthermore, the identification of genes was limited by the availability of references in the databases.

The communities of bark and rhizospheric soil had strong similarities in the abundance of most genes (Supporting Information Fig. S2). The most noticeable difference was the higher abundance of many photosynthetic genes in bark samples, including oxygenic and anoxygenic photosynthesis. The presence of many photosynthetic prokaryotes in bark strongly suggests that these communities have active primary productivity of their own, and do not depend on entirely on consumption of host organic materials. Genes for both oxygenic and anoxygenic photosynthesis were abundant in bark, while they were very rare in the rhizospheric soil samples collected. Although bark surfaces are shaded by trees canopies, enough light might still reach them to sustain significant photosynthetic activity. As many bacteria have pigments which absorb at different wave-lengths of the light spectrum as compared to plants, light reaching the bark surface could be enriched in these wave-lengths, providing enough energy to sustain microbial growth. Measurements of carbon fixation and metabolic flux should be conducted to understand the contribution of this photosynthetic activity to bark microbial communities. The presence of nitrogenase genes in avocado bark suggests a direct input of bioavailable nitrogen to these microbial communities through nitrogen fixation. As these genes belonged mainly to heterocystforming cyanobacteria (Fig. 4), nitrogen fixation could take place under oxic conditions at the bark surface. Nitrogen fixation has been observed on plants surfaces, like leaves, although it is not clear whether the fixed nitrogen can benefit the plants (Lambais *et al*., 2017; Stone *et al*., 2018), for example through run-off from rain water.

Methylotrophy genes were widespread in both rhizospheric soil and bark environments, with different genes for Proteobacteria and Actinobacteria (Fig. 4). However, the absence of methane monooxygenase genes in bark indicates an inability to utilize methane directly. This was unexpected, as significant production of methane by Archaea in the interior of trees stems is known to occur (Yip *et al*., 2019). Perhaps methane diffused through bark is not enough to sustain metanotrophic growth, as most metanotrophs are not facultative, and depend on methane supply for growth. Alternatively, methane oxidation could be performed by Archaea, which were not considered in this analysis. Interestingly, many methylotrophs in bark had also photosynthetic genes. The role of photosynthesis in methylotrophyc bacteria could be as an alternative energy source when one-carbon compounds are not available (Miroshnikov *et al*., 2019). In plants surfaces, these bacteria could benefit from plant-produced sulfide as an electron donor, but this hypothesis needs experimental verification (Atamna-Ismaeel, 2011).

Microorganisms on plant surfaces could be affected by exposure to high light and UV, and it has been shown that pigment synthesis can protect them from photochemical damage (Jacobs *et al*., 2005). Carotenoids have different roles in photosynthetic organisms, acting as antennae pigments, scavenging reactive oxygen and de-exciting chlorophyll under high light and other stresses. Even though the inner stems, including the trunk, can be protected by the canopy, the higher abundance of these genes suggests a higher need of protection form oxidative stress. Similarly, the high abundance of genes for the synthesis of tocopherol, another reactive oxygen scavenger also suggests the need for protection against oxidative stress in the bark environment.

Based of the phyllosphere studies, other adaptations have been proposed to cope with potential environmental stresses on plant surfaces (Vorholt *et al*., 2012). These include exopolysaccharide and surfactant production, quorum sensing, flagellar motility, antimicrobial substance production. Genes for all these functions had many genes detected in both rhizospheric soil and bark (Supporting Information File S2), with few significant differences between them. Although the products of these genes might help combat different stresses, they do not seem to be specific adaptations to plant surface environments.

Many genes for ABC transporters of sugars, sugar alcohols and amino acids were detected, similarly to studies on the phylosphere (Delmotte *et al*., 2009; Lambais *et al*., 2017). This suggests an important presence of heterothrophic bacteria. Consistently, most of these transporters belonged to Actinobacteria, which were one of the most abundant phyla in the bark communities (Aguirre-von-Wobeser *et al*., 2020). The proportion of the organic matter consumed by these bacteria which is plant-derived or produced by photosynthetic microorganisms remains to be determined. A few reads from the bacterial lignin-degraing enzyme called dye-decolorizing peroxidase (de Gonzalo *et al*., 2016) from Actinobacteria were detected in both rhizospheric soil and bark communities, suggesting that some of these bacteria can utilize it.

The present study provides a general picture of the bacterial functions present in bark microbial communities, with emphasis on their difference with rhizospheric soil. The communities have a large photoautothrophic component, which likely provides carbon and energy independently of the trees photosynthesis. Evidence of heterothrophic growth was also observed, which could be sustained by both plant and microbial productivity. Genes for stress adaptation specific to the bark environment were mainly involved in protection from oxidative stress.

To manage bark microbial communities with the introduction of microorganisms of biotechnological interest (for example, Dunlap *et al*., 2017), strains should be selected which could adapt to this environment. Our results suggest different strategies, which could help narrow the search for strains with more probability of establishing stable populations. Phototrophic bacteria could be considered, as they seem to be successful in avocado bark. Alternatively, strains with capacity of utilization of complex organic molecules could benefit from plant and bacteria-derived productivity. Furthermore, strains with ample resistance to animicrobials should be preferred, to avoid growth hindrance from other microorganisms or compounds produced by the plant.

## Supporting information

Supplementary materialas file S1

## Acknowledgments

I thank Ulrike Wiegel Torres, Jesús Torres Rodríguez and José Guadalupe Arellano Lara, owners and caretakers of the studied avocado orchard; Enrique Ibarra Laclette and Emanuel Villafán de la Torre for access to the computer cluster at INECOL and technical support. Financial support for this project was provided by the Mexican Council of Science and Technology (CONACyT) and the Mexican Ministry of Education (SEP), grant CB-2014-01-242956.

The following Supporting Information is available for this article:

**Fig. S1** Comparison of the abundance of all genes detected in all samples of bark and rhizospheric soil of *Persea americana* (Mill.).

**Fig. S2** Statistical tests on the difference in abundance of sequencing reads between bark and rhizospheric soil for each annotated gene.

**Fig. S3** Cyanobacterial photosynthetic antenna genes with different abundance in *Persea americana* (Mill.) bark and rhizospheric soil.

**Fig. S4** Carbon fixation genes with different abundance in *Persea americana* (Mill.) bark and rhizospheric soil.

**Fig. S5** Sucrose and starch genes with different abundance in *Persea americana* (Mill.) bark and rhizospheric soil.

**Table S1** Reads retained after different steps in the analysis.

**Table S2** Enzymes of the Arnon-Buchanan Cycle (Reverse TCA Cycle) in bark and soil.

**Supporting File S1** Abundances of all reads by KEGG annotation and results of statistical tests.

**Supporting File S2** Number of genes identified in soil and bark samples in all prokaryotic KEGG pathways.

## References

Aguirre-von-Wobeser E, Alonso-Sánchez A, Méndez-Bravo A, Villanueva Espino LA, Reverchon F. 2020. Bark from avocado trees of different geographic locations have consistent microbial communities. bioArxiv. doi: doi.org/10.1101/2020.08.21.261396.

Akinpelu DA, Aiyegoro OA, Akinpelu OF, Okoh AI. 2015. Stem bark extract and fraction of *Persea americana* (Mill.) exhibits bactericidal activities against strains of *Bacillus cereus* associated with food poisoning. Molecules 20:416–429.

Antonov IV. 2020. Two cobalt chelatase subunits can be generated from a single *chlD* gene via programmed frameshifting. Molecular Biology and Evolution 37: 2268–2278.

Arrigoni E, Antonielli L, Pindo M, Pertot I, Perazzolli M. 2018. Tissue age and plant genotype affect the microbiota of apple and pear bark. Microbiological Research 211:57–68.

Arrigoni E, Albanese D, Longa CMO, Angeli D, Donati C, Ioriatti C, Pertot I, Perazzolli M. 2020. Tissue age, orchard location and disease management influence the composition of fungal and bacterial communities present on the bark of apple trees. Environmental Microbiology 22:2080–2093.

Atamna-Ismaeel N, Finkel O, Glaser F, von Mering C, Vorholt JA, Koblížek M, Belkin S, Béjà O. 2012. Bacterial anoxygenic photosynthesis on plant leaf surfaces. Environmental Microbiology Reports 4:209–216.

Bolger AM, Lohse M, Usadel B. 2014. Trimmomatic: a flexible trimmer for Illumina sequence data. Bioinformatics 30:2114–2120.

Benjamini Y, Hochberg Y. 1995. Controlling the false discovery rate: a practical and powerful approach to multiple testing. Journal of the Royal Statistical Society Series B 57:289–300.

Buchfink B, Xie C, Huson DH. 2015. Fast and sensitive protein alignment using DIAMOND. Nature Methods 12:59–60.

Cheng C, Wickham JD, Chen L, Xu D, Lu M, Sun J. 2018. Bacterial microbiota protect an invasive bark beetle from a pine defensive compound. Microbiome 6:132.

Coleman-Derr D, Desgarennes D, Fonseca-Garcia C, Gross S, Clingenpeel S, Woyke T, North G, Visel A, Partida-Martinez LP, Tringe SG. 2016. Plant compartment and biogeography affect microbiome composition in cultivated and native *Agave* species. New Phytol 209:798–811.

Crowther TW, Glick HB, Covey KR, Bettigole C, Maynard DS, Thomas SM, Smith JR, Hintler G, Duguid MC, Amatulli G et al. 2015. Mapping tree density at a global scale. Nature 525:201–205.

Dailey HA, Dailey TA, Gerdes S, Jahn D, Jahn M, O’Brian MR, Warren MJ. 2017. Prokaryotic heme biosynthesis: multiple pathways to a common essential product. Microbiology and Molecular Biology Reviews 81:e00048–16.

de Gonzalo G, Colpa DI, Habib MH, Fraaije MW. 2016. Bacterial enzymes involved in lignin degradation. Journal of Biotechnology 236:110–109.

Delmotte N, Knief C, Chaffron S, Innerebner G, Roschitzki B, Schlapbach R, von Mering C, Vorholt, JA. 2009. Community proteogenomics reveals insights into the physiology of phyllosphere bacteria. Proceedings of the National Academy of Sciences 106:16428–16433.

Deschamps AM, Mahoudeau G, Lecelliette L, Lebeault JM. 1980. Isolation and identification of bark decaying and utilizing bacteria of various origins. Revue d’Écologie et de Biologie du Sol 17:577–581.

Deschamps AM, Richard C, Lebeault JM. 1983. Bacteriology and nutrition of environmental strains of Klebsiella pneumoniae involved in wood and bark decay. Annales de l’Institut Pasteur/Microbiologie 134:189–196.

Dunlap CA, Lueschow S, Carrillo D, Rooney AP. 2017. Screening of bacteria for antagonistic activity against phytopathogens of avocados. Plant Gene 11:17–22.

El-Hamalawi ZA, Menge JA. 1994. Avocado trunk canker disease caused by Phytophthora citricola: investigation of factors affecting infection and disease development. Plant Disease 78:260–260.

Erakhrumen AA, Boyi OD. 2016. Micro-organisms present in degrading xylem tissues of some trees and detached parts in University of Benin arboretum, Benin City, Nigeria. Journal of Forestry, Environment and Sustainable Development 2:140–153.

Gorbushina AA. 2007. Life on the rocks. Environmental Microbiology 9:1613–1631.

Gowda DC, Sarathy C, Raju TS. 1988. Structural investigations on the mucilaginous polysaccharides isolated from bark of the avocado tree (*Persea americana* Mill). Carbohydrate Research 15:117–25.

Hacquard S, Schadt CW. 2015. Towards a holistic understanding of the beneficial interactions across the *Populus* microbiome. New Phytologist 205:1424–1430.

Hassani MA, Durán P, Hacquard S. 2018. Microbial interactions within the plant holobiont. Microbiome 6:58.

Jacobs JL, Carroll TL, Sundin GW. 2005. The role of pigmentation, ultraviolet radiation tolerance, and leaf colonization strategies in the epiphytic survival of phyllosphere bacteria. Microbial Ecology 49:104–113.

Kakumanu ML, Cantrell CL, Williams MA. 2013. Microbial community response to varying magnitudes of desiccation in soil: a test of the osmolyte accumulation hypothesis. Soil Biology and Biochemistry 57:644–653.

Kanehisa M, Goto S. 2000. KEGG: kyoto encyclopedia of genes and genomes. Nucleic Acids Research 28:27–30.

Kanehisa M, Sato Y, Morishima K. 2016. BlastKOALA and GhostKOALA: KEGG tools for functional characterization of genome and metagenome sequences. Journal of Molecular Biology 428:726–31.

Kinkar E, Kinkar A, Saleh M. 2019. The multicopper oxidase of *Mycobacterium tuberculosis* (MmcO) exhibits ferroxidase activity and scavenges reactive oxygen species in activated THP-1 cells. International Journal of Medical Microbiology 309:151324.

Lambais MR, Lucheta AR, Crowley DE. 2014. Bacterial community assemblages associated with the phyllosphere, dermosphere, and rhizosphere of tree species of the Atlantic forest are host taxon dependent. Microbial Ecology 68:567–574.

Lambais MR, Barrera SE, Santos EC, Crowley DE, Jumpponen A. 2017. Phyllosphere metaproteomes of trees from the Brazilian Atlantic forest show high levels of functional redundancy. Microbial Ecology 73:123–134.

Langmead B, Trapnell C, Pop M, Salzberg SL. 2009. Ultrafast and memory-efficient alignment of short DNA sequences to the human genome. Genome Biology 10:R25.

Leff JW, Del Tredici P, Friedman WE, Fierer N. 2015. Spatial structuring of bacterial communities within individual *Ginkgo biloba* trees. Environmental Microbiology 17:2352–2361.

Mehltreter K, Flores-Palacios A, García-Franco JG. 2005. Host preferences of low-trunk vascular epiphytes in a cloud forest of Veracruz, Mexico. Journal of Tropical Ecology 21:651–660.

Muñoz-García MA, Melado-Herreros A, Balenzategui JL, Barrerio P. 2014. Low-cost irradiance sensors for irradiation assessments inside tree canopies. Solar Energy 103:143–153.

O’Kennedy BT, Titus JS. 1979. Isolation and mobilization of storage proteins from apple shoot bark. Physiologia Plantarum 45:419–424.

O’Leary NA, Wright MW, Brister JR, Ciufo S, Haddad D, McVeigh R, Rajput B, Robbertse B, Smith-White B, Ako-Adjei D et al. 2016. Reference sequence (RefSeq) database at NCBI: current status, taxonomic expansion, and functional annotation. Nucleic Acids Research 44:D733–D745.

Ottesen AR, Peña AG, White JR, Pettengill JB, Li C, Allard S, Rideout S, Allard M, Hill T, Evans P, Strain E. 2013. Baseline survey of the anatomical microbial ecology of an important food plant: *Solanum lycopersicum* (tomato). BMC Microbiology 13:114.

Roberson EB, Firestone MK. 1992. Relationship between desiccation and exopolysaccharide production in a soil *Pseudomonas* sp. Applied and Environmental Microbiology 58:1284–1291.

Rognes T, Flouri T, Nichols B, Quince C, Mahé F. 2016. VSEARCH: a versatile open source tool for metagenomics. PeerJ 4:e2584.

Rosell JA, Gleason S, Méndez Alonzo R, Chang Y, Westoby M. 2014. Bark functional ecology: evidence for tradeoffs, functional coordination, and environment producing bark diversity. New Phytologist 201:486–497.

Sakuragi Y, Maeda H, DellaPenna D, Bryant DA. 2006. α-Tocopherol plays a role in photosynthesis and macronutrient homeostasis of the cyanobacterium *Synechocystis* sp. PCC 6803 that is independent of its antioxidant function. Plant Physiology 141:508–521.

Selesi D, Schmid M, Hartmann A. 2005. Diversity of green-like and red-like ribulose-1, 5-bisphosphate carboxylase/oxygenase large-subunit genes (cbbL) in differently managed agricultural soils. Applied and Environmental Microbiology 71:175–184.

Sghaier H, Hezbri K, Ghodhbane-Gtari F, Pujic P, Sen A, Daffonchio D, Boudabous A, Tisa LS, Klenk HP, Armengaud J, Normand P. 2016. Stone-dwelling actinobacteria *Blastococcus saxobsidens, Modestobacter marinus* and *Geodermatophilus obscurus* proteogenomes. The ISME Journal 10:21–29.

Stone BW, Weingarten EA, Jackson CR. 2018. The role of the phyllosphere microbiome in plant health and function. Annual Plant Reviews 1:1–24.

Storey JD, Tibshirani R. 2003. Statistical significance for genomewide studies. Proceedings of the National Academy of Sciences 100:9440–9445.

Spangenberg A, Hofmann F, Kirchner M. 2002. Determining the agricultural ammonia immission using bark bio-monitoring: comparison with passive sampler measurements. Journal of Environmental Monitoring 4:865–869.

Tanase C, Coşarcă S, Muntean DL. 2019. A critical review of phenolic compounds extracted from the bark of woody vascular plants and their potential biological activity. Molecules 24:1182.

Vorholt JA. 2012. Microbial life in the phyllosphere. Nature Reviews Microbiology 10:828–840.

Vitulo N, Lemos WJF, Calgaro M, Confalone M, Felis GE, Zapparoli G, Nardi T. 2019. Bark and grape microbiome of *Vitis vinifera*: influence of geographic patterns and agronomic management on bacterial diversity. Frontiers in Microbiology 9:3203.

Wolterbeek HT, Kuik P, Verburg TG, Wamelink GWW, Van Dobben H. 1996. Relations between sulphate, ammonia, nitrate, acidity and trace element concentrations in tree bark in the Netherlands. Environmental Monitoring and Assessment 40:185–201.

Wu Y, Qu M, Pu X, Lin J, Shu B. 2020. Distinct microbial communities among different tissues of citrus tree *Citrus reticulata* cv. Chachiensis. Scientific Reports 10:1–9.

Yatazawa M, Hambali GG, Uchino F. 1983. Nitrogen fixing activity in warty lenticellate tree barks. Soil Science and Plant Nutrition 29:285–294.

Yip DZ, Veach AM, Yang ZK, Cregger MA, Schadt CW. 2019. Methanogenic Archaea dominate mature heartwood habitats of Eastern Cottonwood (*Populus deltoides*). New Phytologist 222:115–121.

