## Supplementary materials Figures and Tables for "Functional metagenomics of bark microbial communities from avocado trees (*Persea americana* Mill.) reveals potential for bacterial primary productivity"

Functional metagenomics of bark microbial communities from avocado trees (*Persea americana* Mill.) reveals widespread oxygenic and anoxygenic prokaryotic photosynthesis genes

Eneas Aguirre-von-Wobeser

### Supporting information

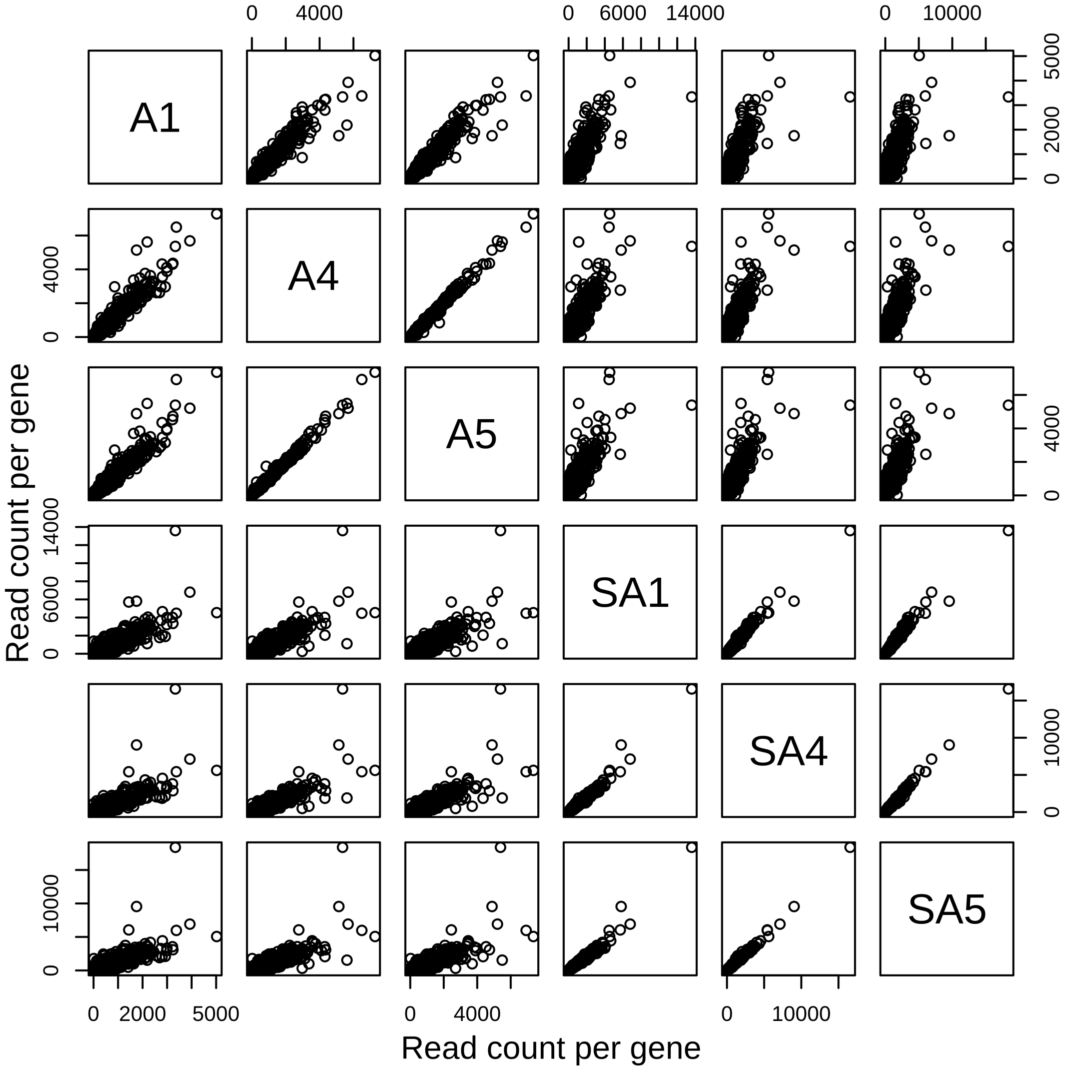


**Fig. S1.** Comparison of the abundance of all genes detected in all samples of bark and rhizospheric soil of Persea americana (Mill.). Samples A1, A4 and A5 correspond to bark, and samples S1, S4 and S5 to rhizospheric soil.

(a)

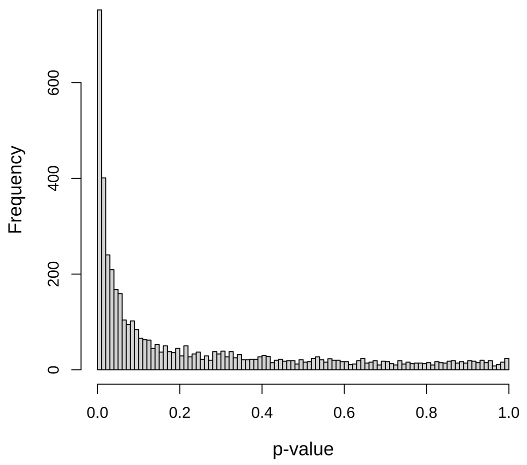


(b)

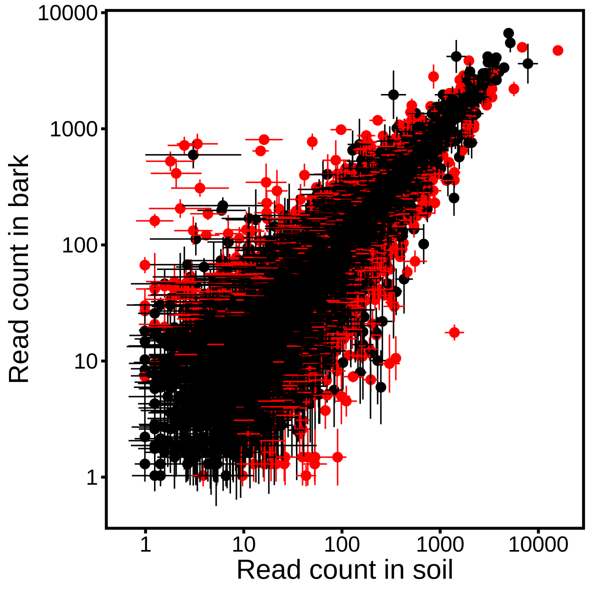


**Fig S2**. Statistical tests on the difference in abundance of sequencing reads between bark and rhizospheric soil for each annotated gene. (a) Distribution of unadjusted p-values from 4425 t-tests conducted, one for each gene, considering genes found in all samples. (b) Genes with significantly different abundance between bark and rhizospheric soil (adjusted $p < 0.05$).

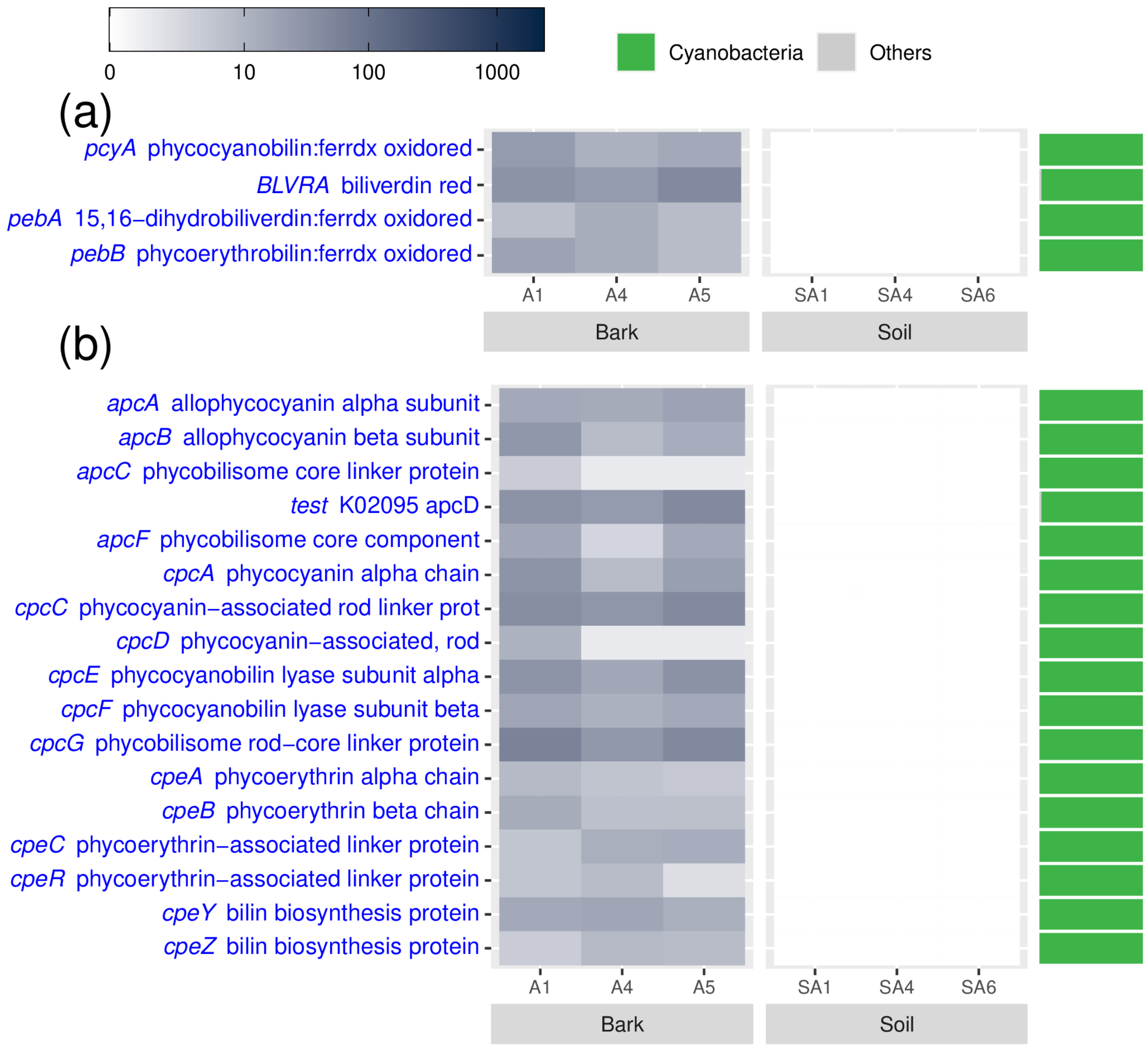


**Fig S3.** Cyanobacterial photosynthetic antenna genes with different abundance in *Persea americana* (Mill.) bark and rhizospheric soil. (a) Phycobin synthesis genes. (b) Biliprotein synthesis genes. Genes shown had significant differences between bark and rhizospheric soil (adjusted $p < 0.05$). The scale shows the number of reads identified for each group, from a randomly selected set of 10,000 reads for each biological sample.

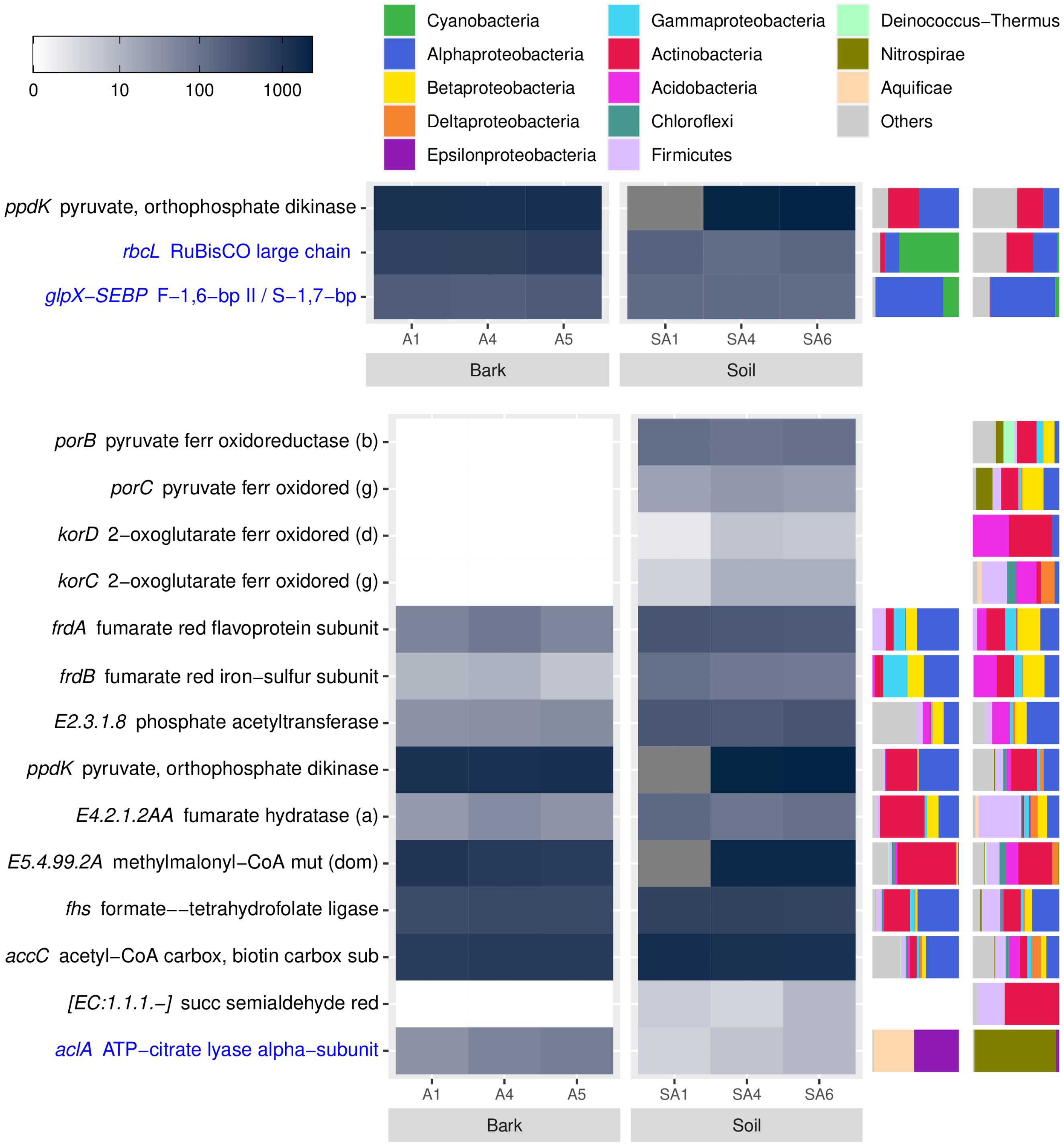


**Fig S4.** Carbon fixation genes with different abundance in *Persea americana* (Mill.) bark and rhizospheric soil. (a) Calvin-Benson cycle genes. (b) Alternative pathways for carbon fixation. Genes shown had significant differences between bark and rhizospheric soil (adjusted $p < 0.05$). Blue font in row labels indicates a higher abundance in bark. The scale shows the number of reads identified for each group, from a randomly

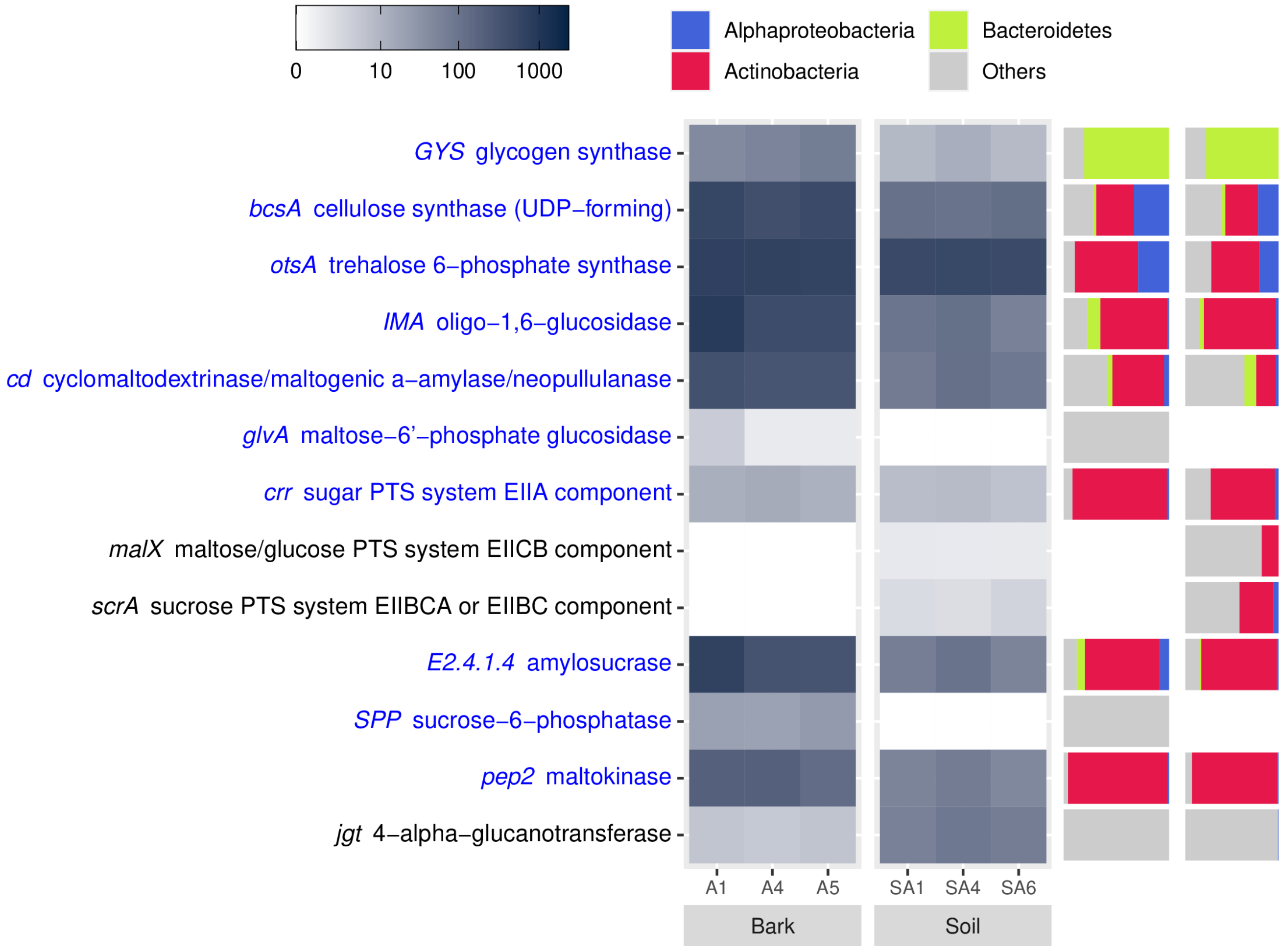


**Fig S5.** Sucrose and starch genes with different abundance in *Persea americana* (Mill.) bark and rhizospheric soil. Genes shown had significant differences between bark and rhizospheric soil (adjusted $p < 0.05$). Blue font in row labels indicates a higher abundance in bark. The scale shows the number of reads identified for each group, from a randomly selected set of 10,000 reads for each biological sample.

**Table S1** Reads retained after different steps in the analysis.

| Sample | Reads after Diamond alignment | Reads after BlastKoala | Reads with KEGG Annotations |
|---|---|---|---|
| A1 | 4010322 | 1879810 (46.9%) | 888905 (22.2%) |
| A4 | 5399996 | 2681880 (49.7%) | 1286098 (23.8%) |
| A5 | 5399994 | 2680907 (49.6%) | 1275946 (23.6%) |
| SA1 | 5317088 | 2487398 (46.8%) | 1135700 (21.4%) |
| SA4 | 5399998 | 2670161 (49.4%) | 1226241 (22.7%) |
| SA5 | 5399998 | 2666346 (49.4%) | 1201409 (22.2%) |

*Note: A sample of 10 million reads was taken before Diamond Alignment.

**Table S2.** Enzymes of the Arnon-Buchanan Cycle (Reverse TCA Cycle) in bark and soil. Values are means of read counts based on subsamples of 10 million reads. Standard deviations are shown in parenthesis.

| Gene | Product name | Bark | Soil |
|---|---|---|---|
| porA | pyruvate ferredoxin oxidoreductase alpha subunit | 0.7(0.6) | 153.3 (48.3) |
| porB | pyruvate ferredoxin oxidoreductase beta subunit | 0 (0) | 110.0 (12.5) |
| porD | pyruvate ferredoxin oxidoreductase delta subunit | 0.3 (0.6) | 40.0 (59.8) |
| porC, porG | pyruvate ferredoxin oxidoreductase gamma subunit | 0 (0) | 18.3 (11.9) |
| por, nifJ | pyruvate-ferredoxin/flavodoxin oxidoreductase | 109.7 (53.9) | 303.0 (416.5) |
| pps, ppsA | pyruvate, water dikinase | 681.7 (33.7) | 266.7 (230.1) |
| ppdK | pyruvate, orthophosphate dikinase | 1269.0 (268.8) | 1430.3 (754.0) |
| ppc | phosphoenolpyruvate carboxylase | 1129.0 (359.6) | 294.7 (416.1) |
| pycA | pyruvate carboxylase subunit A | 1.3 (2.3) | 166.3 (278.6) |
| pycB | pyruvate carboxylase subunit B | 1.7 (1.5) | 404.3 (361.9) |
| PC, pyc | pyruvate carboxylase | 911.0 (86.7) | 824.7 (437.3) |
| mdh | malate dehydrogenase | 753.7 (85.7) | 518.0 (296.7) |
| fumA, fumB | fumarate hydratase, class I | 252.3 (24.9) | 1239.0 (1299.3) |
| fumC, FH | fumarate hydratase, class II | 1128.0 (127.1) | 1797.7 (1938.6) |
| fumA | fumarate hydratase subunit alpha | 33.0 (13.1) | 609.0 (939.9) |
| fumB | fumarate hydratase subunit beta | 9.7 (3.2) | 894.3 (997.3) |
| sdhA, frdA | succinate dehydrogenase / fumarate reductase, flavoprotein subunit | 1975.7 (93.2) | 984.7 (965.1) |
| sdhB, frdB | succinate dehydrogenase / fumarate reductase, iron-sulfur subunit | 731.7 (65.0) | 363.7 (404.4) |
| sdhC, frdC | succinate dehydrogenase / fumarate reductase, cytochrome b subunit | 310.7 (47.7) | 164.0 (123.1) |
| sdhD, frdD | succinate dehydrogenase / fumarate reductase, membrane anchor subunit | 68.7 (22.5) | 23.7 (15.0) |
| frdA | fumarate reductase flavoprotein subunit | 61.0 (23.6) | 149.3 |

| | | | |
|---|---|---|---|
| | | | (88.7) |
| frdB | fumarate reductase iron-sulfur subunit | 8.3 (3.2) | 724.3 (1183.3) |
| frdC | fumarate reductase subunit C | 0.3 (0.6) | 271.7 (461.0) |
| frdD | fumarate reductase subunit D | 1.7 (0.6) | 165.3 (158.0) |
| sucD | succinyl-CoA synthetase alpha subunit | 724.0 (61.7) | 938.3 (458.1) |
| sucC | succinyl-CoA synthetase beta subunit | 971.7 (158.6) | 631.7 (376.9) |
| korA, oorA, oforA | 2-oxoglutarate/2-oxoacid ferredoxin oxidoreductase subunit alpha | 767.0 (88.4) | 971.0 (821.9) |
| korB, oorB, oforB | 2-oxoglutarate/2-oxoacid ferredoxin oxidoreductase subunit beta | 419.0 (38.4) | 1020.7 (373.7) |
| korC, oorC | 2-oxoglutarate ferredoxin oxidoreductase subunit gamma | 0 (0) | 9.7 (4.0) |
| korD, oorD | 2-oxoglutarate ferredoxin oxidoreductase subunit delta | 0 (0) | 252.0 (427.0) |
| IDH1, IDH2, icd | isocitrate dehydrogenase | 1189.0 (59.1) | 529.3 (544.1) |
| ACO, acnA | aconitate hydratase | 2618.0 (310.9) | 1080.3 (1437.0) |
| acnB | aconitate hydratase 2 / 2-methylisocitrate dehydratase | 91.0 (7.8) | 621.7 (1030.0) |
| aclA | ATP-citrate lyase alpha-subunit | 53.0 (24.6) | 59.3 (64.5) |
| aclB | ATP-citrate lyase beta-subunit | 16.3 (9.9) | 7.3 (7.0) |
