## Supplementary materials file S2 for "Functional metagenomics of bark microbial communities from avocado trees (*Persea americana* Mill.) reveals potential for bacterial primary productivity"

**Supporting File 2. Number of genes identified in soil and bark samples in all prokaryotic KEGG pathways.** Counts represent the number of genes present with at least one read in each sample for the corresponding group (soil or bark).

| Carbohydrate metabolism | Soil | Bark |
| --- | --- | --- |
| map00010 Glycolysis / Gluconeogenesis | 77 | 71 |
| map00020 Citrate cycle | 46 | 42 |
| map00030 Pentose phosphate pathway | 58 | 56 |
| map00040 Pentose and glucuronate interconversions | 57 | 57 |
| map00051 Fructose and mannose metabolism | 74 | 74 |
| map00052 Galactose metabolism | 37 | 38 |
| map00053 Ascorbate and aldarate metabolism | 29 | 29 |
| map00500 Starch and sucrose metabolism | 69 | 66 |
| map00520 Amino sugar and nucleotide sugar metabolism | 105 | 102 |
| map00620 Pyruvate metabolism | 80 | 78 |
| map00630 Glyoxylate and dicarboxylate metabolism | 78 | 78 |
| map00640 Propanoate metabolism | 78 | 74 |
| map00650 Butanoate metabolism | 74 | 68 |
| map00660 C5-Branched dibasic acid metabolism | 19 | 17 |
| map00562 Inositol phosphate metabolism | 19 | 19 |
| Energy metabolism |  |  |
| map00190 Oxidative phosphorylation | 89 | 88 |
| map00195 Photosynthesis | 22 | 47 |
| map00196 Photosynthesis - antenna proteins | 2 | 26 |
| map00710 Carbon fixation in photosynthetic organisms | 25 | 25 |
| map00720 Carbon fixation pathways in prokaryotes | 70 | 56 |
| map00680 Methane metabolism | 122 | 103 |
| map00910 Nitrogen metabolism | 45 | 39 |
| map00920 Sulfur metabolism | 75 | 64 |
| Lipid metabolism |  |  |
| map00061 Fatty acid biosynthesis | 24 | 23 |
| map00062 Fatty acid elongation | 2 | 1 |
| map00071 Fatty acid degradation | 26 | 25 |
| map00072 Synthesis and degradation of ketone bodies | 8 | 8 |
| map00073 Cutin, suberine and wax biosynthesis | 1 | 1 |
| map00100 Steroid biosynthesis | 3 | 4 |
| map00561 Glycerolipid metabolism | 31 | 30 |
| map00564 Glycerophospholipid metabolism | 39 | 41 |
| map00565 Ether lipid metabolism | 6 | 7 |
| map00600 Sphingolipid metabolism | 13 | 12 |
| map00590 Arachidonic acid metabolism | 5 | 6 |
| map00591 Linoleic acid metabolism | 2 | 3 |
| map00592 alpha-Linolenic acid metabolism | 4 | 5 |
| map01040 Biosynthesis of unsaturated fatty acids | 9 | 9 |
| Nucleotide metabolism |  |  |
| map00230 Purine metabolism | 115 | 111 |
| map00240 Pyrimidine metabolism | 68 | 65 |
| Amino acid metabolism |  |  |
| map00250 Alanine, aspartate and glutamate metabolism | 44 | 44 |
| map00260 Glycine, serine and threonine metabolism | 78 | 79 |
| map00270 Cysteine and methionine metabolism | 73 | 74 |
| map00280 Valine, leucine and isoleucine degradation | 47 | 44 |
| map00290 Valine, leucine and isoleucine biosynthesis | 15 | 15 |
| map00300 Lysine biosynthesis | 30 | 29 |
| map00310 Lysine degradation | 30 | 30 |
| map00220 Arginine biosynthesis | 40 | 39 |
| map00330 Arginine and proline metabolism | 70 | 65 |
| map00340 Histidine metabolism | 31 | 33 |
| map00350 Tyrosine metabolism | 38 | 39 |
| map00360 Phenylalanine metabolism | 42 | 43 |
| map00380 Tryptophan metabolism | 31 | 31 |
| map00400 Phenylalanine, tyrosine and tryptophan biosynthesis | 46 | 45 |
| Metabolism of other amino acids |  |  |
| map00410 beta-Alanine metabolism | 25 | 25 |
| map00430 Taurine and hypotaurine metabolism | 14 | 14 |
| map00440 Phosphonate and phosphinate metabolism | 16 | 18 |
| map00450 Selenocompound metabolism | 19 | 19 |
| map00460 Cyanoamino acid metabolism | 9 | 9 |
| map00471 D-Glutamine and D-glutamate metabolism | 9 | 9 |
| map00472 D-Arginine and D-ornithine metabolism | 6 | 5 |
| map00473 D-Alanine metabolism | 5 | 5 |
| 00480 Glutathione metabolism | 24 | 23 |
| Glycan biosynthesis and metabolism |  |  |
| map00510 N-Glycan biosynthesis | 3 | 3 |
| map00513 Various types of N-glycan biosynthesis | 3 | 3 |
| map00515 Mannose type O-glycan biosynthesis | 1 | 1 |
| map00514 Other types of O-glycan biosynthesis | 2 | 2 |
| map00531 Glycosaminoglycan degradation | 9 | 9 |
| map00603 Glycosphingolipid biosynthesis - globo and isoglobo series | 3 | 3 |
| map00604 Glycosphingolipid biosynthesis - ganglio series | 1 | 1 |
| map00540 Lipopolysaccharide biosynthesis | 37 | 35 |
| map00550 Peptidoglycan biosynthesis | 34 | 29 |
| map00511 Other glycan degradation | 12 | 12 |
| map00571 Lipoarabinomannan (LAM) biosynthesis | 14 | 12 |
| map00572 Arabinogalactan biosynthesis - Mycobacterium | 10 | 7 |
| Metabolism of cofactors and vitamins |  |  |
| map00730 Thiamine metabolism | 22 | 21 |
| map00740 Riboflavin metabolism | 23 | 16 |
| map00750 Vitamin B6 metabolism | 11 | 10 |
| map00760 Nicotinate and nicotinamide metabolism | 43 | 40 |
| map00770 Pantothenate and CoA biosynthesis | 26 | 27 |
| map00780 Biotin metabolism | 14 | 13 |
| map00785 Lipoic acid metabolism | 5 | 5 |
| map00790 Folate biosynthesis | 43 | 42 |
| map00670 One carbon pool by folate | 22 | 20 |
| map00830 Retinol metabolism | 4 | 4 |
| map00860 Porphyrin and chlorophyll metabolism | 81 | 85 |
| map00130 Ubiquinone and other terpenoid-quinone biosynthesis | 38 | 41 |
| Metabolism of terpenoids and polyketides |  |  |
| map00900 Terpenoid backbone biosynthesis | 29 | 29 |
| map00909 Sesquiterpenoid and triterpenoid biosynthesis | 3 | 3 |
| map00906 Carotenoid biosynthesis | 20 | 21 |
| map00981 Insect hormone biosynthesis | 1 | 1 |
| map00908 Zeatin biosynthesis | 2 | 2 |
| map00903 Limonene and pinene degradation | 7 | 7 |
| map00281 Geraniol degradation | 13 | 13 |
| map01052 Type I polyketide structures | 1 | 1 |
| map01051 Biosynthesis of ansamycins | 4 | 4 |
| map01059 Biosynthesis of enediyne antibiotics | 14 | 17 |
| map01056 Biosynthesis of type II polyketide backbone | 5 | 4 |
| map01057 Biosynthesis of type II polyketide products | 3 | 3 |
| map00253 Tetracycline biosynthesis | 5 | 5 |
| map00523 Polyketide sugar unit biosynthesis | 10 | 10 |
| map01054 Nonribosomal peptide structures | 1 | 1 |
| map01053 Biosynthesis of siderophore group nonribosomal peptides | 13 | 14 |
| map01055 Biosynthesis of vancomycin group antibiotics | 10 | 9 |
| Biosynthesis of other secondary metabolites |  |  |
| map00940 Phenylpropanoid biosynthesis | 7 | 7 |
| map00945 Stilbenoid, diarylheptanoid and gingerol biosynthesis | 1 | 1 |
| map00941 Flavonoid biosynthesis | 1 | 1 |
| map00944 Flavone and flavonol biosynthesis | 1 | 1 |
| map00943 Isoflavonoid biosynthesis | 1 | 1 |
| map00901 Indole alkaloid biosynthesis | 1 | 1 |
| map00950 Isoquinoline alkaloid biosynthesis | 8 | 8 |
| map00960 Tropane, piperidine and pyridine alkaloid biosynthesis | 11 | 11 |
| map00232 Caffeine metabolism | 2 | 2 |
| map00965 Betalain biosynthesis | 3 | 3 |
| map00966 Glucosinolate biosynthesis | 2 | 2 |
| map00311 Penicillin and cephalosporin biosynthesis | 6 | 6 |
| map00332 Carbapenem biosynthesis | 2 | 2 |
| map00261 Monobactam biosynthesis | 15 | 15 |
| map00521 Streptomycin biosynthesis | 12 | 12 |
| map00524 Neomycin, kanamycin and gentamicin biosynthesis | 1 | 1 |
| map00525 Acarbose and validamycin biosynthesis | 5 | 5 |
| map00401 Novobiocin biosynthesis | 8 | 8 |
| map00404 Staurosporine biosynthesis | 2 | 2 |
| map00405 Phenazine biosynthesis | 7 | 6 |
| map00333 Prodigiosin biosynthesis | 3 | 3 |
| Xenobiotics biodegradation and metabolism |  |  |
| map00362 Benzoate degradation | 74 | 71 |
| map00627 Aminobenzoate degradation | 43 | 41 |
| map00364 Fluorobenzoate degradation | 10 | 10 |
| map00625 Chloroalkane and chloroalkene degradation | 15 | 16 |
| map00361 Chlorocyclohexane and chlorobenzene degradation | 17 | 16 |
| map00623 Toluene degradation | 19 | 15 |
| map00622 Xylene degradation | 21 | 23 |
| map00633 Nitrotoluene degradation | 16 | 11 |
| map00642 Ethylbenzene degradation | 5 | 3 |
| map00643 Styrene degradation | 15 | 15 |
| map00791 Atrazine degradation | 10 | 10 |
| map00930 Caprolactam degradation | 12 | 12 |
| map00363 Bisphenol degradation | 2 | 2 |
| map00621 Dioxin degradation | 9 | 9 |
| map00626 Naphthalene degradation | 10 | 10 |
| map00624 Polycyclic aromatic hydrocarbon degradation | 11 | 11 |
| map00365 Furfural degradation | 7 | 7 |
| map00984 Steroid degradation | 12 | 14 |
| map00980 Metabolism of xenobiotics by cytochrome P450 | 3 | 3 |
| map00982 Drug metabolism - cytochrome P450 | 5 | 5 |
| map00983 Drug metabolism - other enzymes | 17 | 17 |
| Transcription |  |  |
| map03020 RNA polymerase | 12 | 12 |
| map03022 Basal transcription factors | 5 | 4 |
| Translation |  |  |
| map03010 Ribosome | 63 | 67 |
| map00970 Aminoacyl-tRNA biosynthesis | 35 | 33 |
| map03013 RNA transport | 8 | 8 |
| map03015 mRNA surveillance pathway | 1 | 2 |
| map03008 Ribosome biogenesis in eukaryotes | 7 | 6 |
| Folding, sorting and degradation |  |  |
| map03060 Protein export | 18 | 18 |
| map04141 Protein processing in endoplasmic reticulum | 5 | 5 |
| map04122 Sulfur relay system | 21 | 18 |
| map03050 Proteasome | 4 | 4 |
| map03018 RNA degradation | 20 | 19 |
| Replication and repair |  |  |
| map03030 DNA replication | 26 | 24 |
| map03410 Base excision repair | 20 | 19 |
| map03420 Nucleotide excision repair | 11 | 10 |
| map03430 Mismatch repair | 23 | 22 |
| map03440 Homologous recombination | 26 | 25 |
| map03450 Non-homologous end-joining | 3 | 2 |
| Membrane transport |  |  |
| map02010 ABC transporters | 357 | 344 |
| map02060 Phosphotransferase system (PTS) | 35 | 30 |
| map03070 Bacterial secretion system | 53 | 54 |
| Signal transduction |  |  |
| map02020 Two-component system | 300 | 262 |
| Transport and catabolism |  |  |
| map04142 Lysosome | 15 | 15 |
| map04146 Peroxisome | 18 | 18 |
| Cell growth and death |  |  |
| map04112 Cell cycle - Caulobacter | 27 | 27 |
| Cellular community - prokaryotes |  |  |
| map02024 Quorum sensing | 110 | 102 |
| map05111 Biofilm formation - Vibrio cholerae | 41 | 37 |
| map02025 Biofilm formation - Pseudomonas aeruginosa | 54 | 51 |
| map02026 Biofilm formation - Escherichia coli | 36 | 30 |
| Cell motility |  |  |
| map02030 Bacterial chemotaxis | 23 | 23 |
| map02040 Flagellar assembly | 37 | 35 |
